# Lysosomal proteolysis of amyloid beta is impeded by fibrils grown in both acidic and neutral pH environments

**DOI:** 10.1101/2021.06.18.449015

**Authors:** Tyler R. Lambeth, Ryan R. Julian

**Affiliations:** Department of Chemistry, University of California, Riverside, California 92521, United States

**Keywords:** amyloid beta, fibrils, cathepsin, lysosome, autophagy

## Abstract

Aggregation of amyloid-beta (Aβ) into extracellular plaques is a well-known hallmark of Alzheimer’s disease (AD). Similarly, autophagic vacuoles, autophagosomes, and other residual bodies within dystrophic neurites, though more difficult to detect, are characteristic features of AD. To explore the potential intersection between these observations, we conducted experiments to assess whether Aβ fibril formation disrupts lysosomal proteolysis. Fibrils constituted from either Aβ 1-40 or Aβ 1-42 were grown under both neutral and acidic pH. The extent of proteolysis by individual cathepsins (L, D, B, and H) was monitored by both thioflavin T fluorescence and liquid-chromatography combined with mass spectrometry. The results show that all Aβ fibrils are resistant to cathepsin digestion, with significant amounts of undigested material remaining for samples of fibrils grown in both neutral and acidic pH. Further analysis revealed that the neutral-grown fibrils are proteolytically resistant throughout the sequence, while the acid-grown fibrils prevented digestion primarily in the C-terminal portion of the sequence. Fibrils grown from Aβ 1-42 are generally more resistant to degradation compared to Aβ 1-40. Overall, the results indicate that Aβ fibrils formed in the neutral pH environments found in intracellular or extracellular spaces may pose the greatest difficulty for complete digestion by the lysosome, particularly when the fibrils are comprised of Aβ 1-42.

## Introduction

Autophagy is a critical process needed to clear cellular waste and free up resources for reuse or energy production. Within this framework, autophagy delivers peptides and proteins to the lysosome where they are digested into constituent amino acids, forming a crucial cog in the gears that drive proteostasis.^1^ Target substrates can be gathered from inside the cell or from the extracellular space through endocytosis.^2^ Due to variety of pathways leading to the lysosome, substrates can be subjected to many conditions and environments prior to fusion with a lysosome. Additionally, the endo/lysosomal system utilizes acidic compartments for delivery and degradation of substrates, further expanding the range of different environments that may be experienced prior to degradation.^3^ Failure of the endo/lysosomal system can lead to a variety of complications, including a class of diseases known as lysosomal storage disorders. Lysosomal storage is most frequently caused by hydrolase dysfunction, which leads to the accumulation of undigested substrates and eventual failure of the organelle.^4^ Autophagic disruption has also been implicated in Alzheimer’s disease (AD) due to the hallmark observation of lysosomal storage.^5,6^

Inside the lysosome, hydrolases known as cathepsins degrade peptides and proteins into the constituent amino acids which are then released by transporter proteins back into the cytosol.^7^ While a few members of the cathepsin family are exopeptidases which cleave from the termini, the majority are endopeptidases which cleave somewhere in the middle of the sequence.^8^ Among the endopeptidases, the most abundant and active are cathepsin L (catL, all cathepsins will be abbreviated similarly) and catD.^9^ Studies have demonstrated that knocking out either catD or catL induces pathology and death in mice within 4 weeks.^10,11^ The most abundant lysosomal exopeptidase, catB, recognizes and binds to the C-terminus (as well as the C-terminal mimic L-isoAsp)^12^ and removes two amino acids at a time as dipeptides. The complementary aminopeptidase, catH, works from the N-terminus to remove one amino acid at a time. Although catB and catH are primarily exopeptidases, they both possess a secondary endopeptidase activity.^13,14^ Knockouts of catB also cause shortened lifespans^10^, while knockouts of catH lead to significantly reduced levels of important neurotransmitter peptides.^15^ These studies demonstrate that while the cathepsins perform similar functions, each enzyme is individually vital to maintain proteostasis.

Amyloid beta (Aβ) is known to aggregate into fibrils that are a prevalent pathological feature of AD.^16^ As target of autophagy, Aβ (including fibrils) can be trafficked from both intracellular and extracellular spaces to the lysosome.^17^ Similar to other amyloid proteins which self-assemble into aggregates such as human islet amyloid polypeptide^18^ and CsgA^19^, Aβ is able to spontaneously form fibril structures in solution via side-chain and backbone interactions. Aβ fibrils are comprised primarily of stacked beta-sheet structures stabilized by hydrophobic interactions along the middle and C-terminal region of the sequence.^20,21^ Amyloid secondary structures have been shown to impede digestion by proteases such as trypsin^22^ and bovine brain proteases^23^ which easily digest monomeric Aβ. In a related system, α-synuclein aggregates were found to resist degradation by catL.^24^ These previous experiments suggest that many amyloid fibrils are resistant in varying degrees to degradation by proteases.

In the case of Aβ, the pH during fibril formation can also alter the nature of the fibril that is formed. For example, fibrils formed in acidic environments differ from those formed at neutral pH. Importantly, the resulting fibrils do not interconvert if the pH is shifted after fibril have formed.^25^ The structural difference between the fibrils is attributed to varying protonation states of histidine residues in the sequence.^26^ While other ionizable groups are unaffected by a shift from cellular to lysosomal pH, the pKa value for the imidazole group comprising the sidechain of histidine lies in the middle of the relevant pH range. As a consequence, histidine is likely to become protonated in acidic compartments. The formation of amyloid beta fibrils is driven by hydrophobic interactions between the peptide chains, and as such can potentially be altered by hydrophilic charges.^27^ By substituting alanine for histidine residues in Aβ40, it has been demonstrated that protonation of sidechains for His6, His13, and His14 significantly affects fibril structure by disfavoring amyloid sheets in the N-terminal half of the sequence.^28^ Due to the origination of fibrils in neutral and acidic cellular spaces, fibrils with varied secondary structures could be delivered to the lysosome for degradation.

Herein, we examine incubations of fibrils grown at cellular and acidic pH with lysosomal cathepsins to evaluate their ability to degrade these structures. Analysis was performed via thioflavin T (ThT) fluorescence to measure the general extent of degradation, and proteolytic products were also quantitatively assessed with liquid-chromatography/mass spectrometry. Significant differences were observed in the amount of degradation as a function of the pH used to grow the fibrils and whether the fibrils were composed of Aβ40 or Aβ42. Further examination of these digestions with mass spectrometry revealed differences in both the identified sequences and length of peptides remaining after incubation. By plotting these products as a function of intensity, we were able to map proteolytically resistant regions as a function of fibril composition and pH during formation.

## Experimental Procedures

### Fibril Formation

Lyophilized Aβ powder was purchased from Anaspec. The samples were disentangled via ammonium hydroxide treatment.^29^ 100 µg aliquots of each peptide were dissolved in 50 µL of 1% ammonium hydroxide solution (w/v) with sonication followed by dilution to 1 mL with either 50 mM tris pH 7.2 or acetate pH 5 buffer for fibril growth. Amyloid beta 1-42 aliquots were fibrilized at 25 µM while amyloid beta 1-40 were fibrilized at 100 µM to start fibril growth after brief agitation. Fibrils were grown for 5 days at 37°C and checked by ThT fluorescence to confirm fibril presence.

### Cathepsin Incubations

Aliquots containing Aβ were digested by cathepsins in acetate buffer pH 5, with 1 mM ethylenediaminetetraacetic acid (EDTA) and 500 µM dithiothreitol (DTT) to prevent active site oxidation. For each digestion 0.4 µg of enzyme were incubated with 20 µg of amyloid beta for a 1:50 enzyme:substrate ratio (w/w). A control sample was set up for each digestion with no enzyme added. Incubations occurred over an 18hour period at 37°C. Digestions were quenched by dilution with 200 mM tris before immediate fluorescent measurements.

### Fluorescence Measurements

The presence of beta-sheet rich aggregates was examined by ThT assay. Samples were diluted to 2 µM in 200 mM tris buffer with 6 µM ThT. Emission scans were performed on a QuantaMaster-400 fluorimeter using an excitation wavelength of 440 nm and an emission wavelength of 485 nm.

### LC-MS Analysis

Samples were analyzed on a Thermo Fisher Ultimate 3000 RSLCnano System interfaced with a Thermo Fisher Velos Pro Orbitrap using an electrospray ionization (ESI) source. Peptides were separated on a capillary column packed in-house with C18 3 µm resin using a Shotgun Proteomics Inc high pressure vessel. Mobile phase A was water 0.1% formic acid and mobile phase B was 80% acetonitrile in water with 0.1% formic acid. Nano-ESI was performed using a spray voltage of 2.1 kV with an S-lens value of 65.

## Results and Discussion

Amyloid beta 1-40 (Aβ40) and 1-42 (Aβ42) were incubated at pH 5 and pH 7.2 to produce stable fibrils with different structures as discussed in the introduction. These fibrils were then digested over an 18-hour period in separate experiments by cathepsin L, D, B, and H, which represent two crucial endopeptidases, one carboxypeptidase, and one aminopeptidase. The extent of digestion was coarsely measured by ThT fluorescence intensity, as shown in Fig. 1a for the digestion of neutral Aβ42 by catL. ThT is a fluorescent dye used to measure the presence of protein and peptide aggregates due to its ability to fluoresce intensely when bound to beta-sheet rich structures.^30^ The fractional intensity of fluorescence remaining after digestion represents the amount of remaining beta-sheet rich structures present in the sample after proteolysis. The fluorescence data from all cathepsin digestions is compiled in Fig 1b. For the endopeptidases, catL digested more fibril relative to catD. For the acid-grown fibrils (in green), catL reduced the amount of fluorescence by >80% from the initial level. However, for the neutral-grown fibrils less digestion was observed, particularly for Aβ42, which only exhibited a ∼30% reduction in fluorescence intensity. Similarly, catD reduced ThT fluorescence less for neutral-grown fibrils less, and yielded almost no change for the neutral Aβ42 fibrils.

**Figure 1.**
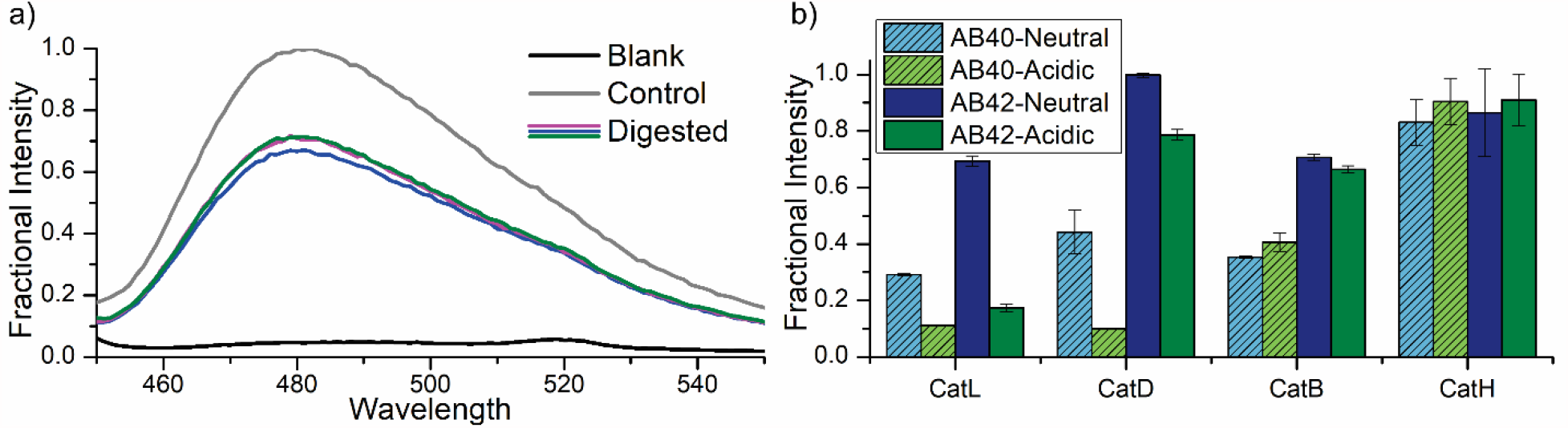
(a) Three replicates for ThT fluorescence following digestion of Aβ42-Neutral by catL. (b) Compiled fluorescence data for all cathepsin incubations.

Endopeptidases operate by binding to several amino acids on either side of the peptide bond targeted for hydrolysis, and as such are potentially sensitive to differences in substrate backbone structure in either direction of the surrounding sequence region. In contrast, catB and catH act primarily as exopeptidases and have the strongest interactions with residues to one side of the targeted peptide bond. For catH, digestion of Aβ42 and Aβ40 yielded similar results and reductions in ThT fluorescence were not significant for either acidic or neutral fibrils. This suggests that catH was unable to access the amyloid region lending ThT fluorescence, although it is unclear whether any portion of the N-terminus was digested. For catB, which attacks from the C-terminal side, significantly more ThT fluorescence was retained by the Aβ42 fibrils for both acidic and neutral fibers. Indeed, neutral or acidic had little effect on the catB results, suggesting that differences in fibril structure likely occur near the N-terminus (as discussed in the introduction). Although there are clearly differences in the ThT results for our various test conditions, more detailed information would likely facilitate greater understanding.

To more precisely determine the outcome of each cathepsin digestion, the samples were analyzed with a combination of liquid-chromatography and mass spectrometry. These experiments were able to identify the precise peptides remaining following proteolysis and defibrilization. Raw chromatograms of the digestion of Aβ42 with catL are shown in Fig. 2a and 2b for neutral and acidic fibrils, respectively. Notable differences in retention times and relative intensities are apparent, suggesting that the two experiments generated significantly different peptide profiles. This possibility is confirmed by MS analysis, which is illustrated schematically in Fig. 2c. Each identified sequence from the chromatograms in Fig. 2a/b are displayed as individual lines in Fig. 2c, where the length of each line maps out the corresponding peptide sequence in relation to the full sequence shown in the middle of the diagram. The sequences are also ordered by intensity, with more intense peptides displayed closer to the full Aβ42 sequence. The full list of peptides can also be found in Tables S1 and S2 in the supporting information. Peptides located in the C-terminal portion of Aβ42 were found for both acidic and neutral fibrils, indicating resistance to catL digestion in this region. However, for the neutral fibrils, a variety of peptides were found from the N-terminal region, including many peptides that contained the N-terminus itself. These results suggest that the amyloid forming region is resistant to digestion for both acidic and neutral fibrils, but that the N-terminal region is accessible and susceptible to digestion for acidic fibrils. The data from Fig. 2c can be more succinctly summarized with additional analysis as shown in Fig. 2d and 2e. In Fig. 2d, histograms of residual peptide length are shown for four different experiments: neutral Aβ40, acidic Aβ40, neutral Aβ42, and acidic Aβ42. Aβ42 digestion products skew more towards the longer sequence lengths in general while Aβ40 digestion products are clustered in smaller length peptides. Notably, the 26-30 length bin contains only intensity from neutral-grown fibrils of Aβ40 and Aβ42. When compared to the catL digestion of monomeric Aβ42 collected in Table S3 and displayed as histograms in Fig. S1, a striking difference is observed. In the monomeric digest, nearly 80% of the peptide products are of lengths less than 5 amino acids long, and there are no peptides observed greater than 15 residues long. This contrast in peptide lengths demonstrates a significant obstruction is occurring in the proteolysis of Aβ fibrils.

**Figure 2.**
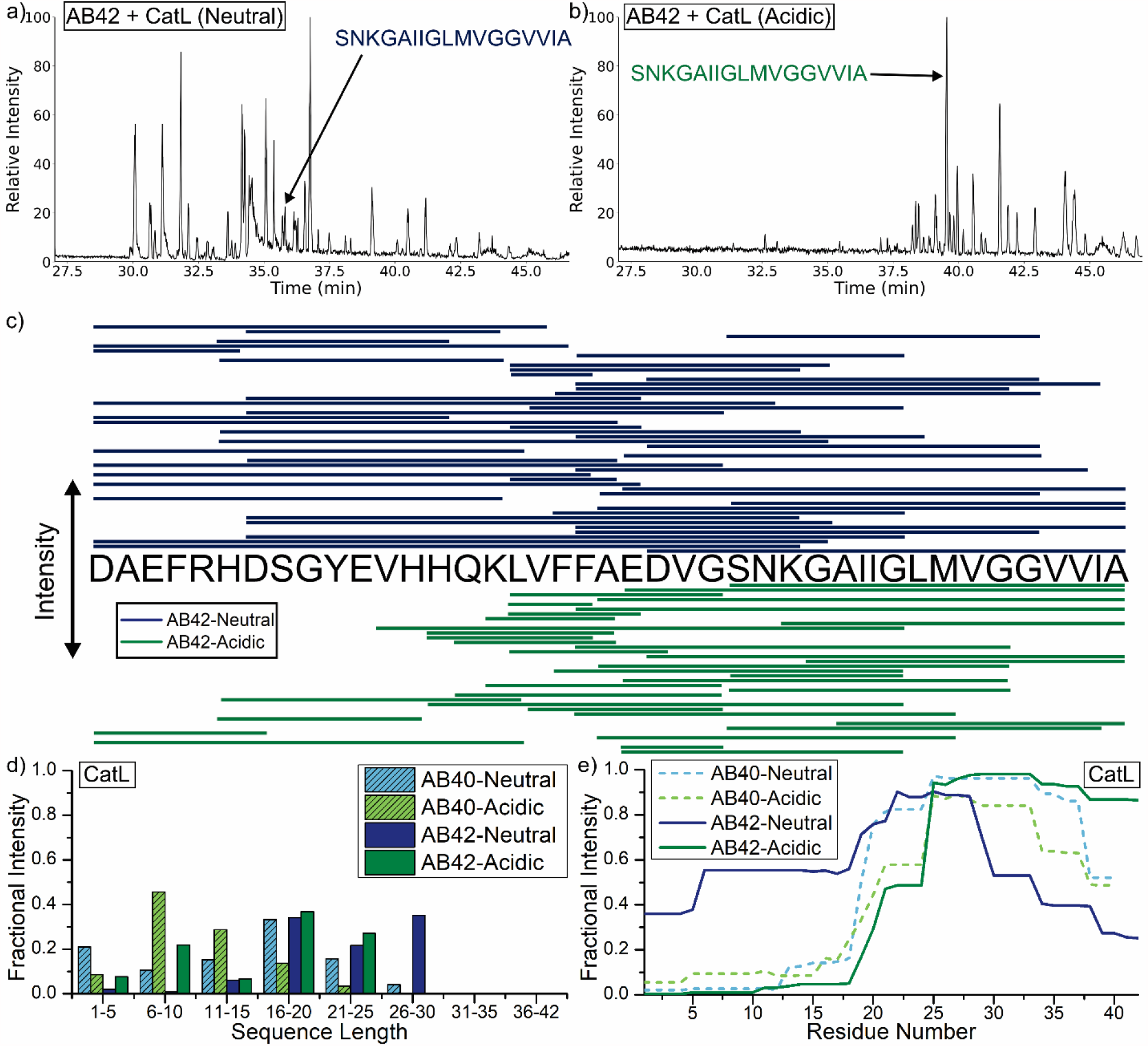
(a) Raw chromatogram for the digestion of Aβ42-neutral and (b) Aβ42-acidic by catL. The proteolytic product Aβ 16-42 is shown in both chromatograms as a reference. (c) Compiled line diagram of the identified peptides from the chromatograms. Lines indicate which part of the full sequence comprises the proteolytic product. Products are ordered by intensity, with the most intense products listed closer to the full sequence. (d) Bar plot showing the length of the identified product peptides. (e) Plot of the residue intensity among the total intensity of all peptides with a length greater than 10, representing areas resistant to proteolysis.

To visually display a summary of the regions with the most resistance to proteolysis, sequences with a length of at least 11 amino acids were used to generate an intensity map as shown in Fig. 2e. The fractional intensity value (y-axis) for each residue (x-axis) was calculated by adding up the intensity of all peptides containing the residue and then dividing by the intensity of all peptides longer than 11 residues. Accordingly, a fractional intensity value of 0.5 means that the residue is present in 50% of the total peptide intensity. Examination of the solid lines (derived from Aβ42 data) reveals a quantitative assessment in excellent agreement with the semi-quantitative representation shown in Fig. 2c. Again, differences in product profiles between various digestions are apparent. The C-terminal region is resistant to proteolysis in all cases. Interestingly, these same fibrils are the most resistant in the N-terminal region. For Aβ40, less difference is noted between digestion of acidic and neutral fibrils. These results are consistent with the fluorescence data shown in Fig. 1b, where the neutral-grown fibrils display a higher amount of intensity after digestion in both Aβ40 and Aβ42, demonstrating an increase in proteolytic resistance.

The same analysis was performed on digestions with the endopeptidase catD, as shown in Fig. 3. The peptide lengths shown in Fig. 3a consist of a tighter spread than those seen with catL digestions. Interestingly, the neutral-grown Aβ42 products are almost entirely sequences with lengths of 16-20 residues. Although most of the intensity from Aβ40 digestions is found in the 16-20 length as well, some peptides were also identified with lengths greater than 20. The absence of shorter peptides may suggest that fewer acceptable binding sites exist for catD or that it is less capable of digesting fibrils in general. The catD digestion of monomeric Aβ42 is illustrated in Table S4 and displayed in Fig. S2. The distribution of intensities suggests that catD is able to more easily bind and degrade sequences in the monomer form, populating the 6-10 and 11-15 bins with around 65% of the product intensity. Additionally, the 1-5 length bin contains a small portion of the intensity, while none is present in either Aβ42 fibril digestion. The peptide intensity map shown in Fig. 3b reveals some similar trends to the previous catL digestions. All digestions showed a resistant region in the C-terminal half of the sequence, extending from around Lys16 to Met35. In the neutral-grown Aβ42, nearly half of the peptides originating from the N-terminal half of the sequence survive digestion. The Aβ40 digests for both acidic and neutral fibrils also yielded more N-terminal peptide intensity. The fluorescence data for these samples shown in Fig. 1b is higher for both digestions of neutral-grown Aβ, which indicates that the resistant residues in the N-terminal half may be contributing to the formation of greater amounts of stacked beta-sheet content involved in ThT binding.

**Figure 3.**
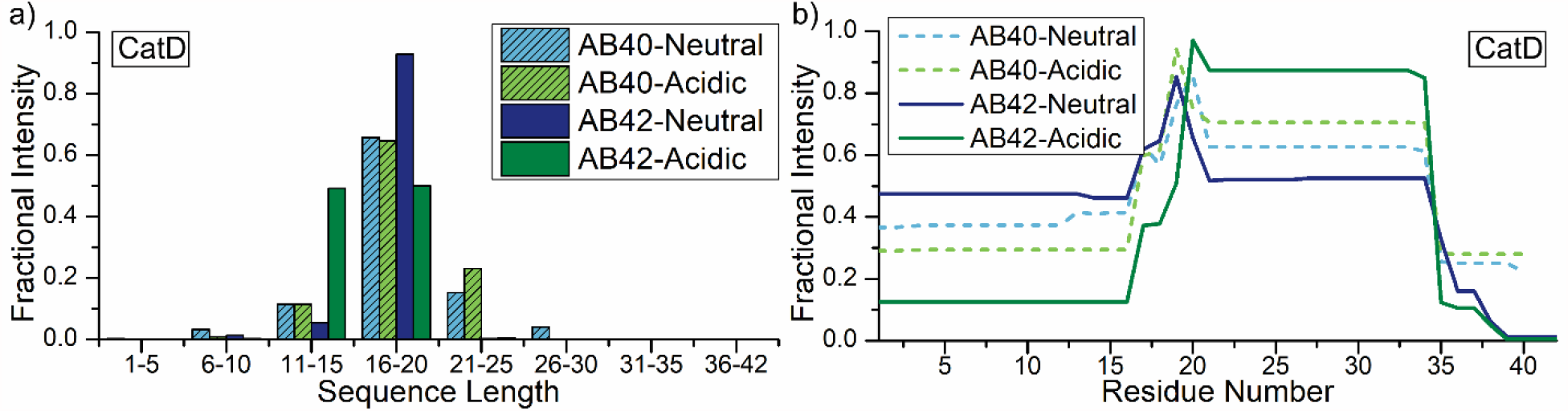
(a) Bar plot of the sequence length of identified proteolytic products from catD incubations. (b) Plot of the residue intensity among the total intensity of all peptides with a length greater than 10.

Identical analyses for results from catB and catH are shown in Fig. 4. These enzymes act primarily as exopeptidases, though both possess some endopeptidase activity. Examining the data for the catB digests in Fig. 4a reveals a wide spread of peptide lengths. The Aβ42 neutral fibril distribution is skewed towards longer lengths, while the Aβ40 neutral fibrils are distributed more towards the center of the distribution. Acidic fibrils from both Aβs populate bimodal distributions, favoring longer and shorter peptides. The peptide intensity map is shown in Fig. 4b. Although catB is a carboxypeptidase which cleaves from the C-terminus, it is unable to progress very far before encountering resistance. Indeed, only the acidic fibrils from Aβ40 reveal any cleavage at the C-terminus that is nearly complete. Ironically, most of the degradation for catB takes place due to secondary endopeptidase activity in the N-terminal region.

**Figure 4.**
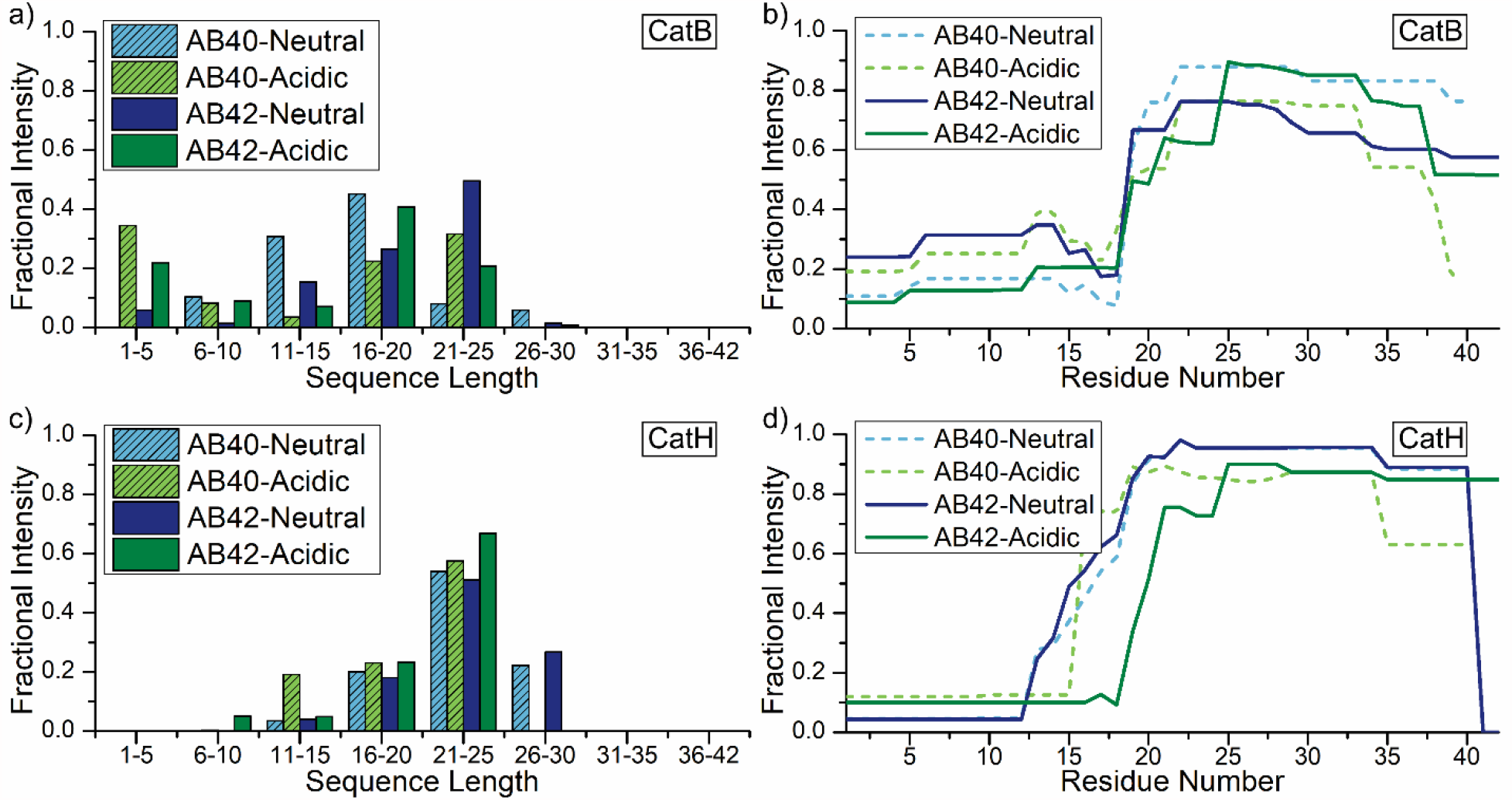
(a) Bar plot of the sequence length of identified proteolytic products from exopeptidase catB and (c) catH incubations. (b) Plot of the residue intensity among the total intensity of all peptides with a length greater than 10 for catB incubations and (d) catH incubations.

In the length histograms for catH shown in Fig. 4c, the majority of all peptide intensity is observed in peptides of length 21-25. CatH is an aminopeptidase that preferentially cleaves one amino acid at a time and is likely to produce products that will not be retained by LC. Notably, peptides of the longest observed length (26-30) were only recorded for the neutral-grown fibrils. In the residue intensity plot for catH shown in Fig. 4d, the N-terminal region is similarly digested for all four experiments. The primary differences in Fig. 4d relate to Aβ42. The acidic fibrils allow greater penetration in the N-terminal region, while an unexpected cleavage at the C-terminus is observed for the neutral fibrils. This C-terminal cleavage goes essentially to completion and is more efficient than the C-terminal cleavage observed for catB. The smaller differences between these digestion profiles are similarly reflected in the smaller differences between the fluorescence intensity shown in Fig. 1b.

## Conclusion

We have examined the influence of fibril formation on the proteolysis of both Aβ40 and Aβ42 in a detailed and quantitative fashion. It is clear that fibril formation interferes with proteolysis by the major lysosomal cathepsins in every case. The differential digestion obtained for fibrils formed at either neutral or acidic pH confirms the likelihood that such fibrils have distinct structures, mostly related to the N-terminal portion of the sequence. Overall, our results suggest that fibrils composed of Aβ42 and formed at neutral pH will present the greatest difficulty for digestion within the lysosome. In contrast, monomeric Aβ is easily digested by cathepsins and appears unlikely to contribute to lysosomal pathology. However, our results suggest that it is possible for amyloid fibrils contribute to AD pathology and the lysosomal storage observed in the disease by simply avoiding degradation. Accumulation of undigested fibrils within lysosomal compartments would eventually disrupt both autophagy and eventually proteostasis, sending affected cells on a pathway towards apoptosis.

## Acknowledgements

The authors gratefully acknowledge funding from the National Institute on Aging, R01AG066626.

## References

1 Kaushik, S., Cuervo, A.M. Proteostasis and Aging. Nat. Med. 2015, 21(12), 1406–1415.

2 Mizushima, N. Autophagy: Process and Function. Genes Dev. 2007, 21(22), 2861–2873.

3 Luzio, J.P., Hackmann, Y., Dieckmann, N.M.G., Griffiths, G.M. The Biogenesis of Lysosomes and Lysosome-Related Organelles. Cold Spring Harbor Perspect. Biol. 2014, 6(9), a016840–a016840.

4 Kiselyov, K., Jennings, J.J., Jr., Rbaibi, Y., Chu, C.T. Autophagy, Mitochondria, and Cell Death in Lysosomal Storage Diseases. Autophagy 2007, 3(3), 259–262.

5 Wolfe, D.M., Nixon, R.A. Autophagy Failure in Alzheimer’s Disease and Lysosomal Storage Disorders: A Common Pathway To Neurodegeneration? Autophagy of the Nervous System; World Scientific, 2018; pp 237–257.

6 Lambeth, T.R., Riggs, D.L., Talbert, L.E., Tang, J., Coburn, E., Kang, A.S., Noll, J., Augello, C., Ford, B.D., Julian, R.R. Spontaneous Isomerization of Long-Lived Proteins Provides a Molecular Mechanism for the Lysosomal Failure Observed in Alzheimer’s Disease. ACS Cent. Sci. 2019, 5, 1387–1395.

7 Bissa, B., Beedle, A., Govindarajan, R. Lysosomal Solute Carrier Transporters Gain Momentum in Research. Clin. Pharmacol. Ther. 2016, 100(5), 431–436.

8 Turk, V., Stoka, V., Vasiljeva, O., Renko, M., Sun, T., Turk, B., Turk, D. Cysteine cathepsins: From structure, function and regulation to new frontiers. Biochim. Biophys. Acta 2012, 1824, 68–88.

9 Turk, B., Turk, D., Turk, V. Lysosomal cysteine proteases: More than scavengers. Biochim. Biophys. Acta. 2000, 1477, 98–111

10 Felbor, U., Kessler, B., Mothes, W., Goebels, H.H., Ploegh, H.L., Bronson, R.T., Olsen, B.R. Neuronal Loss and Brain Atrophy in Mice Lacking Cathepsin B and L. Proc. Natl. Acad. Sci. U.S.A. 2002, 99(12), 7883–7888.

11 Koike, M. Cathepsin D Deficiency Induces Lysosomal Storage with Ceroid Lipofuscin in Mouse CNS Neurons. Neurosci. Res. 2000, 38(18), S29.

12 Lambeth, T.R., Dai, Z., Zhang, Y., Julian, R.R. A two-trick pony: Lysosomal protease cathepsin B possesses surprising ligase activity. RSC Chem. Biol. 2021, 2, 606–611.

13 Quraishi, I., Nägler, D.K., Fox, T., Sivaraman, J., Cygler, M., Mort, J.S., Storer, A.C. The occluding loop in cathepsin B defines the pH dependence of inhibition by its propetide. Biochemistry 1999, 38(16), 5017–5023.

14 Kirschke, H., Langner, J., Wideranders, B., Ansorge, S., Bohley, P., Hanson, H. Acta. Biol. Med. Ger. 1977, 36, 185–199.

15 Lu, W.D., Funkelstein, L., Toneff, T., Reinheckel, T., Peters, C., Hook, V. Cathepsin H functions as an aminopeptidase in secretory vesicles for production of enkephalin and galanin peptide neurotransmitters. J. Neurochem. 2012, 122, 512–522.

16 Ghiso, J., Frangione, B. Amyloidosis and Alzheimer’s disease. Adv. Drug Delivery Rev. 2002, 54(12), 1539–1551.

17 Ciechanover, A., Kwon, Y.T Degradation of misfolded proteins in neurodegenerative diseases: therapeutic targets and strategies. Exp. Mol. Med. 2015, 47, e147.

18 Lam, Y.P.Y., Wootton, C.A., Hands-Portman, I., Wei, J., Chiu, C.K.C., Romero-Canelon, I., Lermyte, F., Barrow, M.P., O’Connor, P.B. Determination of the Aggregate Binding Site of Amyloid Protofibrils Using Electron Capture Dissociation Tandem Mass Spectrometry. J. Am. Soc. Mass Spectrom. 2020, 31(2), 267–276.

19 Wang, H., Shu, Q., Rempel, D.L., Frieden, C., Gross, M.L. Understanding curli amyloid-protein aggregation by hydrogen-deuterium exchange and mass spectrometry. Int. J. Mass Spectrom. 2016, 420, 16–23.

20 Colvin, M.T., Silvers, R., Ni, Q.Z., Can, T.V., Sergeyev, I., Rosay, M., Donovan, K.J., Michael, B., Wall, J., Linse, S., Griffin, R.G. Atomic Resolution Structure of Monomorphic Aβ42 Amyloid Fibrils. J. Am. Chem. Soc. 2016, 138, 9663–9674.

21 Zhang, Y., Rempel, D.L., Zhang, J., Sharma, A.K., Mirica, L.M., Gross, M.L. Pulsed hydrogen-deuterium exchange mass spectrometry probes conformational changes in amyloid beta (Aβ) peptide aggregation. PNAS, 2013, 110(36), 14604–14609.

22 Kheterpal, I., Williams, A., Murphy, C., Bledsoe, B., Wetzel, R. Structural Features of the Aβ Amyloid Fibril Elucidated by Limited Proteolysis. Biochemistry 2001, 40(39), 11757–11767.

23 Chauhan, V., Sheikh, A.M., Chauhan, A., Spivack, W.D., Fenko, M.D., Malik, M.N. Fibrillar amyloid beta-protein inhibits the activity of high molecular weight brain protease and trypsin. J.Alzheimers Dis. 2005, 7(1), 37–44.

24 McGlinchey, R.P., Dominah, G.A., Lee, J.C. Taking a Bite Out of Amyloid: Mechanistic Insights into α-Synuclein Degradation by Cathepsin L. Biochemistry 2017, 56(30), 3881–3884.

25 Wood, S.J., Maleeff, B., Hart, T., Wetzel, R. Physical, Morphological, and Functional Differences between pH 5.8 and 7.4 Aggregates of the Alzheimer’s Amyloid Peptide Aβ. J. Mol. Biol. 1996, 256, 870–877.

26 Shi, H., Li, H., Gong, W., Gong, R., Qian, A., Lee, J.Y., Guo, W. Structural and Binding Properties on Aβ Mature Fibrils Due to the Histidine Tautomeric Effect. ACS Chem. Neurosci. 2019, 10, 4612–4618.

27 Paravatsu, A.K., Petkova, A.T., Tycko, R. Polymorphic Fibril Formation by Residues 10-40 of the Alzheimer’s ß-Amyloid Peptide. Biophys. J. 2006, 90, 4618–4629.

28 Brännsträm, K., Islam, T., Sandblad, L., Olofsson, A. The role of histidines in amyloid ß fibril assembly. FEBS Lett. 2017, 591, 1167–1175.

29 Ryan, T.M., Caine, J., Mertens, H.D.T., Kirby, N., Nigro, J., Breheney, K., Waddington, L.J., Streltsov, V.A., Curtain, C., Masters, C.L. Roberts, B.R. Ammonium hydroxide treatment of A beta produces an aggregate free solution suitable for biophysical and cell culture characterization. PeerJ. 1:e73, 2013.

30 Biancalana, M., Koide, S. Molecular Mechanism of Thioflavin-T binding to amyloid fibrils. Biochim. Biophys. Acta, Proteins and Proteomics. 2010, 1804(7), 1405–1412.

